# Disruption of the Lacunar Canalicular Network in Type 2 Diabetes: Impaired Osteocyte Connectivity in Zucker Diabetic Rats

**DOI:** 10.1101/2025.10.29.683583

**Authors:** Michael Sieverts, Attila Gyulassy, Ishita Juluru, Valerio Pascucci, Claire Acevedo

**Affiliations:** Department of Mechanical Engineering, University of Utah, 1495 E 100 S, Salt Lake City, UT, 84112, USA; Scientific Computing and Imaging Institute, University of Utah, 72 Central Campus Dr, Salt Lake City, UT 84112; Kahlert School of Computing, University of Utah, 1495 E 100 S, Salt Lake City, UT, 84112, USA; Department of Mechanical and Aerospace Engineering, University of California San Diego, San Diego, CA, 92161, USA

## Abstract

Type 2 diabetes affects multiple organ systems, including the skeletal system. Diabetes reduces bone’s mechanical properties and impacts bone cells, such as osteocytes, which are crucial to preserving bone health. Osteocytes maintain bone health through the lacunar canalicular network (LCN), a highly interconnected system vital for remodeling, mechanotransduction, and nutrient transport. Yet the specific impact diabetes has on this network has remained unclear. Here, we used confocal laser scanning microscopy combined with advanced connectomics modeling to achieve high-resolution, three-dimensional reconstructions of the LCN in Zucker Diabetic Sprague Dawley rats, a polygenic model that closely mimics human type 2 diabetes. Diabetes profoundly disrupted LCN connectivity in the femoral mid-cortex, with canalicular and node density reduced by 21% and 30%, respectively. Additionally, we observed a 30–40% increase in lacunar density and highly connected nodes. These architectural shifts impair bone permeability, diminishing mechanosensitivity and compromising nutrient and oxygen transport. Our findings uncover a previously unrecognized mechanism of skeletal fragility in diabetes and highlight the LCN as a promising therapeutic target.

## Introduction

Type 2 diabetes has emerged as a global epidemic with broad consequences in multiple organ systems [1]. Beyond its well-established effects on renal, cardiovascular, and nervous systems, diabetes also has significant impacts on the skeletal system [2, 3]. These effects include altered mechanical properties, mineralization disturbances, and increased microdamage [4]. As a result, patients with diabetes are at an elevated risk of fracture [2–6]. As the prevalence of diabetes continues to rise [7], it is increasingly important to understanding the intricate mechanisms by which it compromises bone quality and microstructure.

The mechanisms underlying elevated fracture risk in patients with diabetes remain an active area of research [2, 3]. Patients with type 2 diabetes often have normal to elevated bone mineral density (BMD) but still experience an increased fracture risk [2, 8]. Some factors contributing to this fragility include the accumulation of advanced glycation end-products (AGEs) at the collagen level and increased cortical porosity at the microstructural level [2, 4, 9–11]. Additionally, type 2 diabetes is believed to alter the differentiation and function of bone cells, particularly osteocytes [2, 11].

Osteocytes are the most abundant bone cells and are essential in maintaining skeletal homeostasis [12]. These cells are housed in small voids within the bone matrix called lacunae, and they extend dendritic processes through small tunnels called canaliculi. Osteocytes connect and communicate through an intricate porous network of lacunae and canaliculi, which comprises the lacunar canalicular network (LCN). This expansive network exhibits an impressive density of approximately 74 kilometers per cubic centimeter in healthy human osteons [13]. Through the LCN, osteocytes regulate bone quality by facilitating essential functions such as cellular communication, remodeling, mechanotransduction, and nutrient transport [12, 14]. Various conditions cause disruptons to this network, such as aging [15] and osteoporosis [16]. Disruptions to this network could reduce bone quality and increase fracture risk [14].

Evidence from high-fat diet mouse models suggest diabetes alters osteocyte morphology without changing the number of canaliculi or dendrite processes [17, 18]. However, limited research has focused on how diabetes affects the LCN in polygenic animal models that more closely mimic human type 2 diabetes. In prior work, our lab demonstrated that type 2 diabetes in a polygenic rat model reduces whole-bone mechanical properties and alters bone microstructure [19]. These findings raise the possibility that the osteocyte LCN could be compromised by diabetes. We hypothesize that diabetes decreases the canalicular density within the LCN, impairing osteocyte function and reducing bone quality. This hypothesis is further supported by data linking diabetes to reduced osteocyte viability and altered LCN connectivity in humans [11, 20, 21], as well as parallels with aging where declines in canalicular density and osteocyte function are linked to increased bone fragility [22–24].

Connectomic analysis techniques are a powerful approach to characterize the LCN [25]. This approach represents the LCN as a topological graph of nodes (junctions of canaliculi) and edges (canaliculi themselves) [25]. Connectomics has been used to identify the deterioration of the LCN with age [26], and connectivity changes in high-fat mouse models [17]. To our knowledge, connectomics has not been used to investigate how type 2 diabetes impacts the LCN in polygenic animal models, such as the Zucker Diabetic Sprague Dawley (ZDSD).

In this study, we combined CLSM with connectomic analysis to quantify how type 2 diabetes alters the LCN in (ZDSD) rats [27]. We report significant reductions in canalicular and node density in the femoral mid-cortex, leading to impaired connectivity and reduced bone permeability. This impact is observed in mid-cortex of the bone and not in regions near the endosteum. These findings establish the LCN as a critical site of diabetes-induced skeletal deterioration and provide new insights into the mechanisms of bone fragility in diabetes.

## Results

### The LCN in the mid-cortex and the endosteum have distinct organizations

To inspect the impacts of diabetes on the LCN, we acquired CLSM images of bone samples from Sprague Dawley rats (Control, N=6) and Zucker Diabetic Sprague Dawley (ZDSD) rats (Diabetic, N=7). To visualize and quantify the characteristics of the LCN, we imaged each bone sample in the mid-cortex and near the endosteum because these regions have distinct organization and structure [28, 29]. We imaged all the bones in the anterior medial quadrant of the femur, where we would expect the bone to be in compression. Features within these samples were segmented using a combination of deep learning semantic segmentation and manual correction (Figure 1a–b).

**Figure 1:**
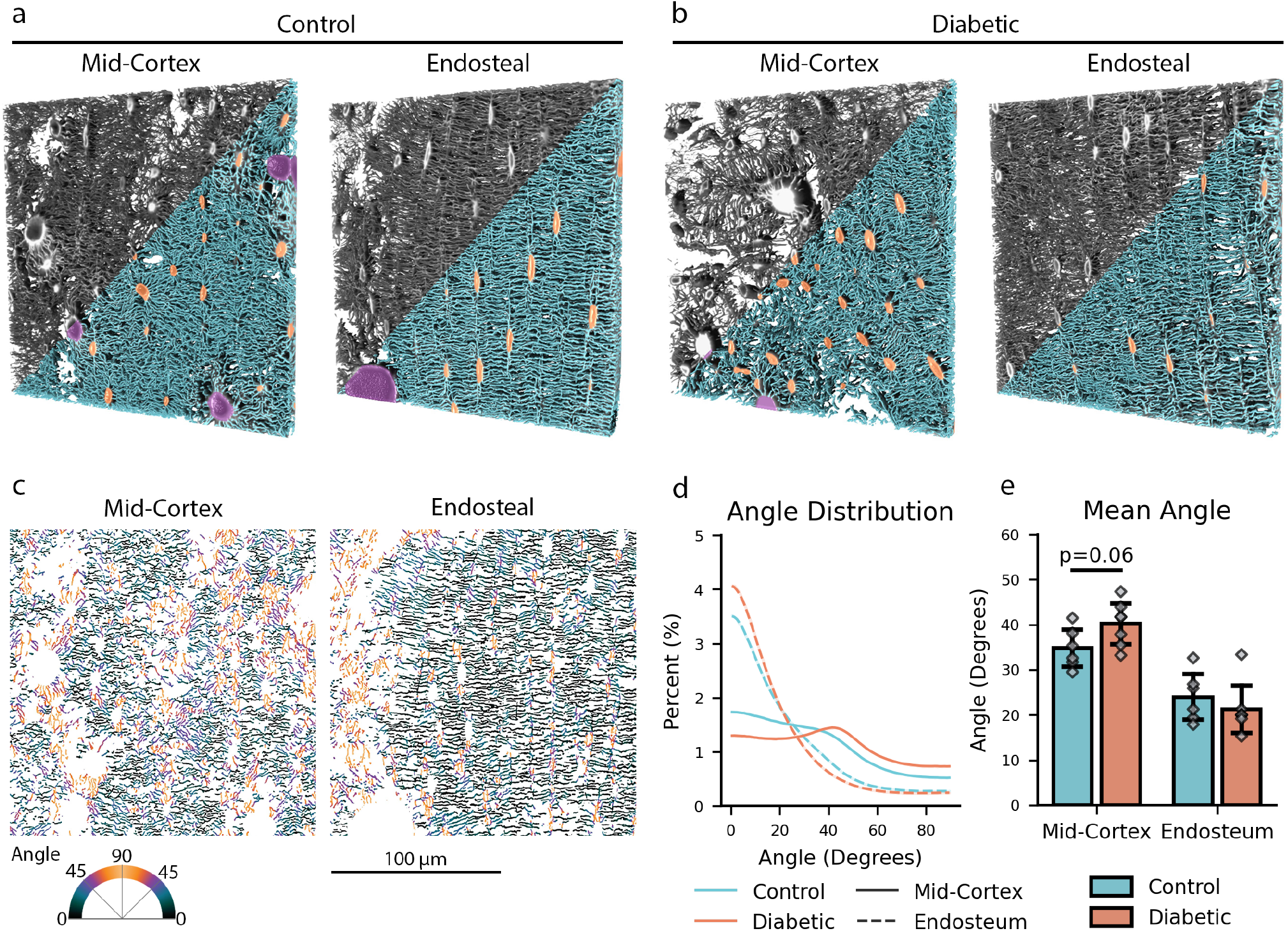
The LCN has a distinct organization in the mid-cortex and the endosteum in both the control and diabetic bone. (**a–b**) 3D reconstructions of 184×184×10 µm volumes imaged using confocal microscopy of both control and diabetic bone. The segmentation of the LCN is shown in these volumes, with the canaliculi labeled in blue, the lacunae labeled in orange, and the blood vessels labeled in purple. (**c**) Example 2D orientation maps of the control bone in both the mid-cortex and the endosteum. The angles range from 0 degrees (horizontal) to 90 degrees (vertical). The LCN is aligned in the endosteal bone, while the LCN is more radially organized in the mid-cortex. (**d**) Average angle distribution of the canaliculi for all samples. In the endosteal bone, most canaliculi are aligned horizontally, near a 0-degree angle. In the mid-cortex, there is a more uniform distribution of angles between 0–50 degrees. (**e**) Bar plot showing the mean angle for each group. The mean angle is generally higher in the mid-cortex, suggesting a more radially organized LCN. All bar graphs show the mean and standard deviation for each measurement. Note that a thin slab was selected for visualization in this figure. Analysis was performed on the 184×184×42 µm volumes.

We measured the 2D orientation angle of the canaliculi to interrogate organizational differences between the two regions and experimental groups. Similar to previous research, we observed that the LCN near the endosteum is highly regular and aligned in a primary direction, while the LCN in the mid-cortex is less regularly aligned (Figure 1c) [28, 29]. In the endosteum, we found that 80% of the canaliculi are oriented between angles 0–37 degrees. In the mid-cortex, 80% of the canaliculi are oriented between angles 0–60 degrees (Figure 1d). These results confirm that the LCN near the endosteum is highly aligned and that most canaliculi in this region are oriented in the direction of the thickness of the bone (horizontally). In contrast, in the mid-cortex, the canaliculi have a more uniform distribution of angles between 0–50 degrees, resulting in a more radial arrangement (Figure 1d). We observe this regional organizational difference in samples from the control and the diabetic groups. Generally, the canalicular orientation depended more on region than the treatment group; however, we also found a 16% (p=0.06) increase in mean angle in the diabetic group in the mid-cortex compared to the control, suggesting an increase in radial orientation in this region due to diabetes (Figure 1e). These observations confirm the heterogeneity of the LCN and show that properties of the LCN vary in different regions of the bone. Additionally, we observe that this heterogeneity persists in diabetic bone.

### Diabetic bone has reduced canalicular network density in the mid-cortex

To perform a connectomic analysis of the LCN, we converted the segmented images to a topological graph of nodes and edges. The nodes are junctions where three or more canaliculi meet, and the edges consist of the canaliculi that link the nodes [25]. To understand the impact that diabetes has on the LCN, we measured properties related to network density for all samples [25]. We quantified canalicular density, the total length of canaliculi per unit volume, and node density, the number of nodes per unit volume [25]. We also measured the lacunar density, the number of lacunae per unit volume, and the mean lacunar volume. These metrics help describe the health and function of the LCN.

In diabetic bone, the LCN network density is significantly reduced in the mid-cortex and unchanged near the endosteum. In the mid-cortex, we observe a 21% (p=0.012) reduction in canalicular density and a 30% (p=0.010) reduction in node density due to diabetes (Figure 3a–b). With the reduction in node density, we also observed an increase in link length in the mid-cortex due to diabetes (Figure 3e). A slight increase, 19% (p=0.070), in canalicular spacing further supports this reduction in network density (Figure 3d). These reductions in network density are only observed in the mid-cortex of the diabetic bones. The density of the LCN in the endosteal region of the diabetic bones was unchanged compared to the controls (Figure 3a–b). These results suggest that diabetes significantly impacts the LCN by reducing the network density. These results indicate that diabetes impacts the LCN in the mid-cortex but not near the endosteum.

**Figure 2:**
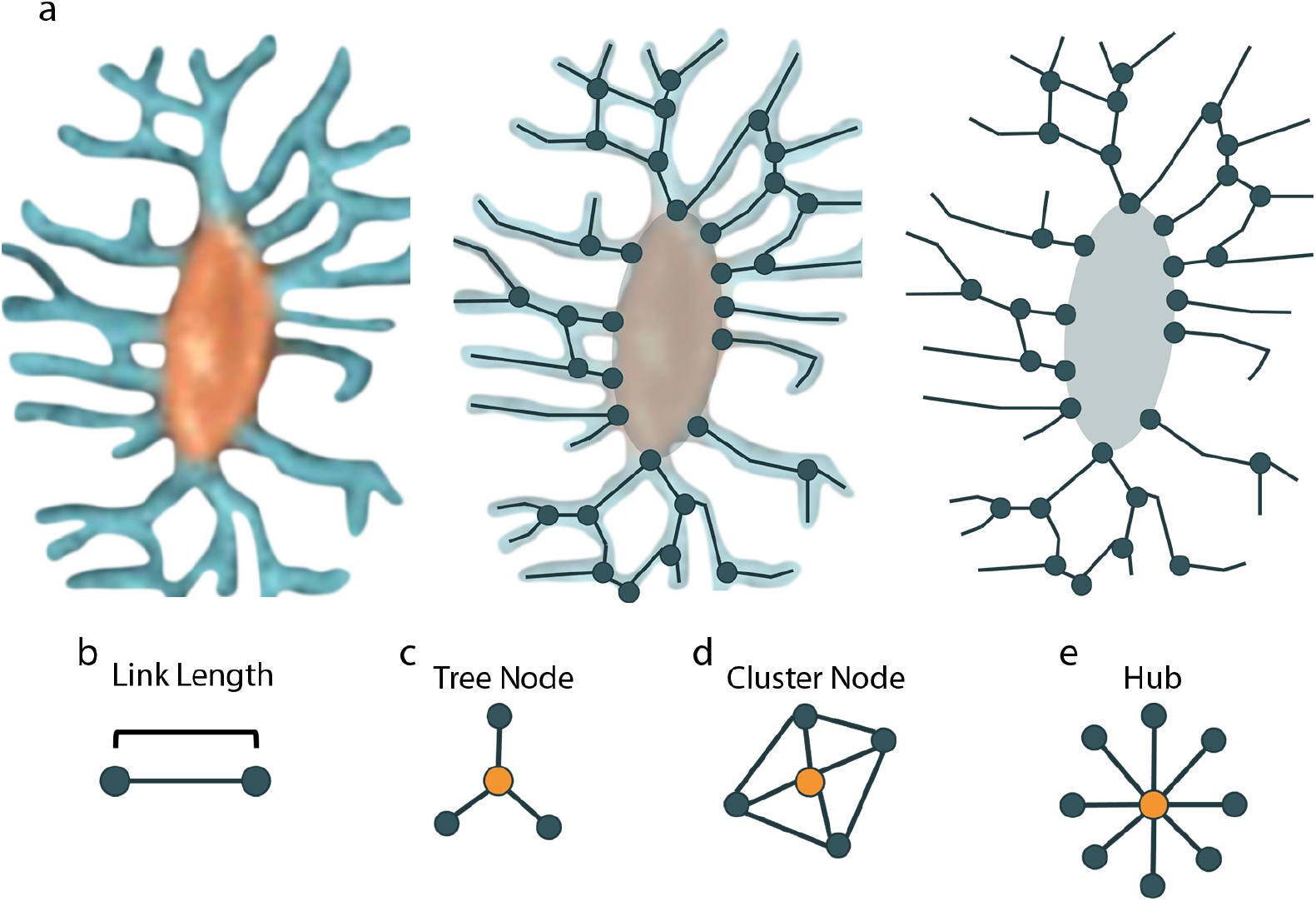
Overview of the LCN being represented as a topological graph. (**a**) An example of how the segmented LCN is converted to a graph consisting of nodes and edges. The lacuna is segmented in orange, and the canaliculi are segmented in blue. Together, these microstructural features constitute the LCN. (**b–e**) Examples of network properties quantified in this research. In these descriptions, the node of interest is highlighted in orange. (**b**) Link length measures the distance between nodes. (**c**) Tree nodes are defined by having a degree of 3 and a clustering coefficient (CC) of 0. (**d**) Cluster nodes are defined by having a CC greater than 0.5. These nodes can have any degree. (**e**) Hubs are defined by having a degree greater than or equal to 8. These nodes can have any CC. Other researchers have used these metrics to describe aspects of the LCN [17, 25, 26, 30].

**Figure 3:**
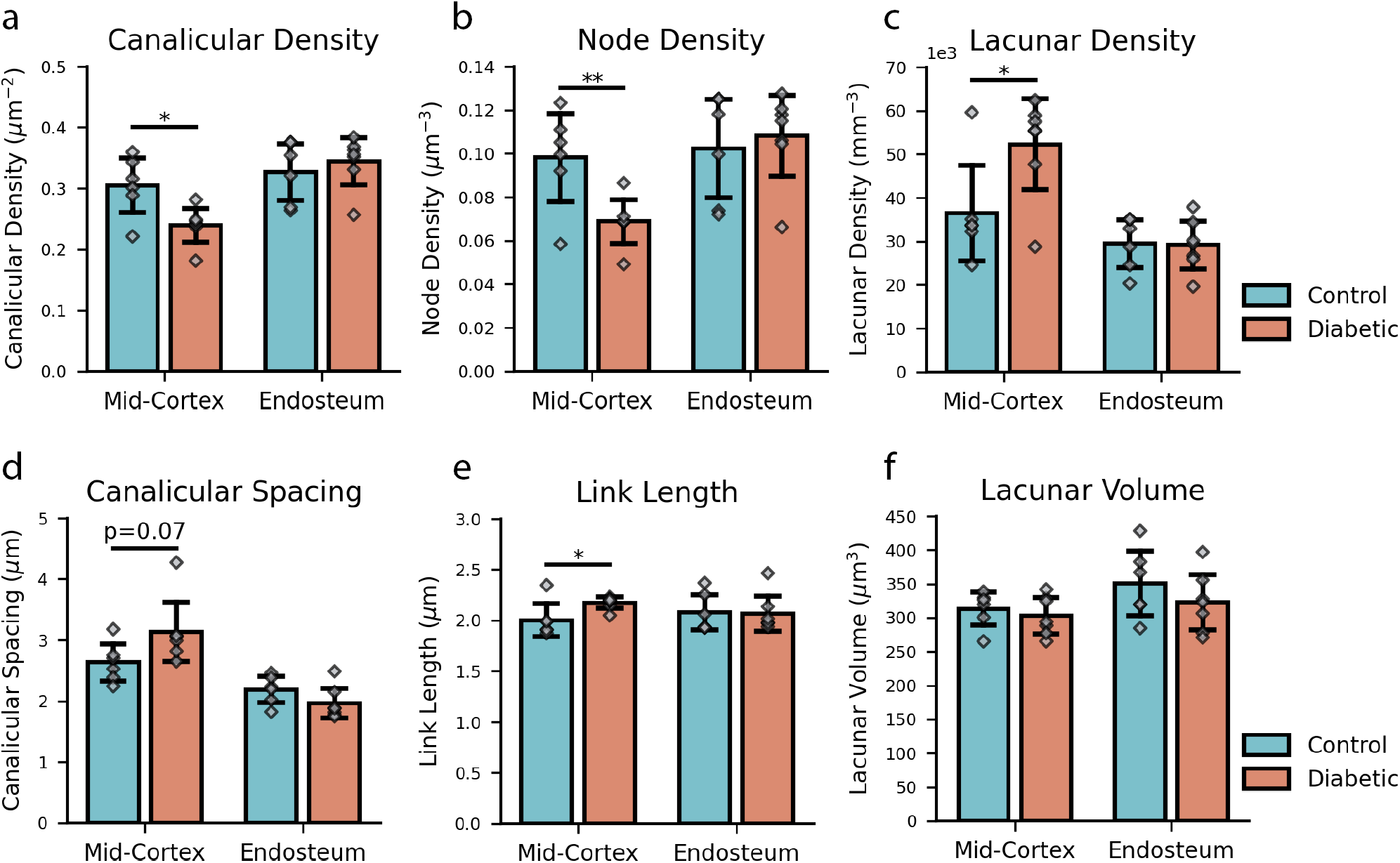
The LCN in diabetic bone has reduced network density in the mid-cortex. (**a–c**) Bar graphs for canalicular, node, and lacunar density. These metrics describe the amount of the LCN in the samples. In the mid-cortex, diabetes results in significant reductions in canalicular density (p=0.012) and node density (p=0.010), as well as a significant increase in lacunar density (p=0.030). (**d–f**) Bar graphs for canalicular spacing, link length, and lacunar volume. In the mid-cortex, the increase in canalicular spacing and link length helps describe the reductions in canalicular and node density. Decreased canalicular density results in increased canalicular spacing. The decreased node density results in increased link length or space between nodes. All bar graphs show the mean and standard deviation for each measurement.

While other properties of the LCN are decreased in the mid-cortex due to diabetes, we observe an increase in lacunar density. We found a significant 43% (p=0.03) increase in lacunar density in the diabetic group in the mid-cortex (Figure 3c), and no change in lacunar volume (Figure 3f). These results show that diabetes has different effects on the canalicular and the lacunar portion of the LCN.

### There is reduced LCN permeability in the mid-cortex of diabetic bone

The LCN graph was represented as a pore-network model (PNM) to evaluate changes in permeability. Using this representation of the LCN, we quantified permeability in three directions: radially, from the endosteum toward the periosteum; circumferentially, around the circumference of the bone; and longitudinally, through the long axis of the bone. We measured a normalized conductance for the LCN in each of these directions. In the radial direction, in the mid-cortex, we observed a significant 41% (p=0.021) decrease in permeability in the diabetic group (Figure 4a). This change in permeability was also visualized in Figure 4d. This visualization demonstrates the reduction in canalicular density and the associated impact on permeability (Figure 4d). In the endosteum, we found a slight 15% increase in permeability with diabetes (Figure 4a). In the circumferential direction, in the mid-cortex, we found a moderate 28% decrease in permeability in the diabetic group (Figure 4b). In the endosteum, there was almost no difference due to diabetes (Figure 4b). In the longitudinal direction, all groups were very similar for both locations within the bone (Figure 4c).

**Figure 4:**
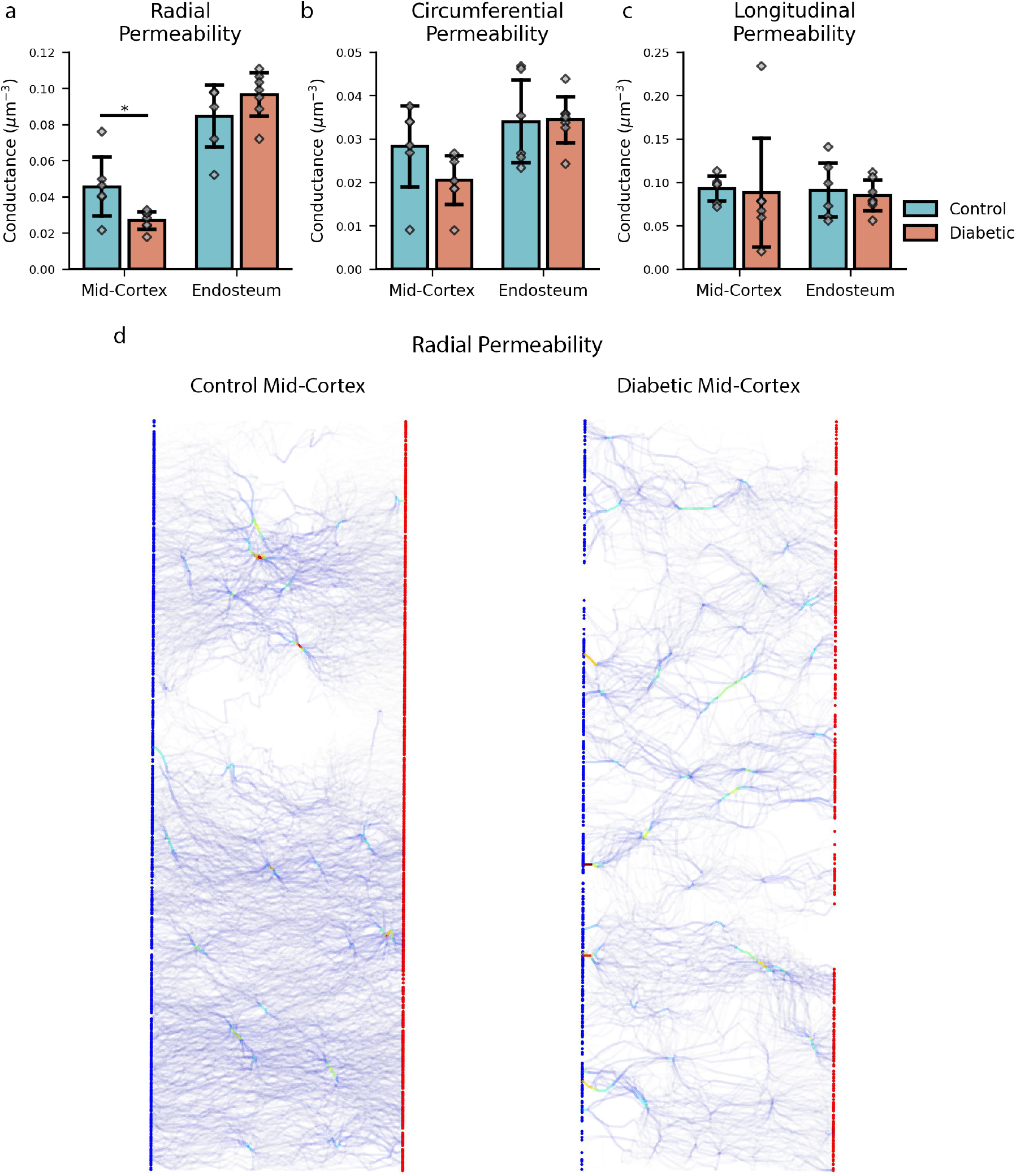
Reduced LCN permeability in the mid-cortex of diabetic bone. (**a**) A bar graph showing the radial permeability of the LCN. In the mid-cortex, diabetes results in a significant 41% reduction in permeability (p=0.021) (**b**) A bar graph showing the circumferential permeability of the LCN. There is still a trend of 28% reduction in permeability (nonsignificant). (**c**) A bar graph showing the longitudinal permeability of the LCN. No change in permeability was found. (**d**) Visualization of the radial permeability of control and diabetic bones in the mid-cortex. The red nodes represent sources on the endosteal side of the bone, and the blue nodes represent sinks on the periosteal side of the bone. All bar graphs show the mean and standard deviation for each measurement.

### The bone matrix is further from the LCN in the mid-cortex of diabetic bone

To investigate changes in how the bone matrix is situated about the LCN, we analyzed the distance to the network for the bone matrix (Figure 5). In the mid-cortex, the diabetic bone has significantly less of the bone matrix located in regions 0.1–0.9 µm compared to the control (Figure 5b). The bone matrix in the mid-cortex of the diabetic bone is 18% (p=0.08) further from the LCN compared to the control (Figure 5c). These results indicate that diabetes causes the bone matrix to be less accessible by the LCN in the mid-cortex of cortical bone.

**Figure 5:**
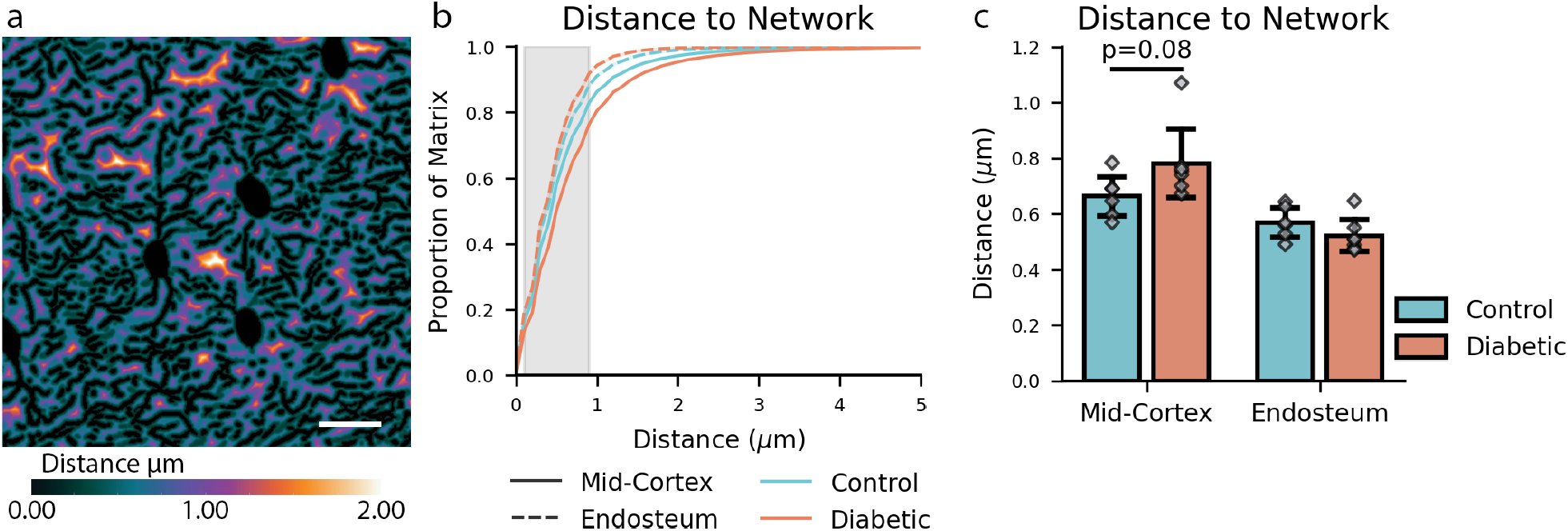
The bone matrix in diabetic bone is further from the LCN in the mid-cortex. (**a**) Representative image of a 3D distance map computed away from the LCN. The scalebar is 10 µm. (**b**) A line plot shows the proportion of bone matrix included at certain distances away from the LCN. In the mid-cortex, the diabetic bone has significantly less of the bone matrix located in the highlighted region between 0.1–0.9 µm. (**c**) A bar plot shows each group’s mean distance to the bone matrix from the LCN. All bar graphs show the mean and standard deviation for each measurement.

### There is an elevated proportion of high-degree nodes in the mid-cortex of diabetic bone

We took an approach similar to Kollmannsberger et al. [30] to evaluate the different types of nodes within the LCN. We classified nodes based on their degree and clustering coefficient (CC). Node degree describes how many edges connect at that point. The node’s CC is a local connectivity measurement [30]. Nodes with a degree of 3 and a CC of 0 were considered tree nodes. These nodes efficiently minimize the impact of local damage, such as a microcrack, on the network’s communication [17]. Nodes of any degree with a CC greater than 0.5 were considered cluster nodes. These nodes have high local interconnectivity [30]. We also evaluated the proportion of hubs within the bone. A node was determined to be a hub if it had a degree of 8 or greater. These high-degree nodes are less resilient to local disruptions. If a hub is compromised, it may disrupt a large portion of the LCN [17]. We found that most of the nodes (≈ 75%) in the LCN are tree nodes for both the control and diabetic groups, regardless of location in the bone (Figure 6a). We observed a low proportion of cluster nodes within the network for all groups; however, there is a 30% (p=0.15) increase in cluster nodes in the diabetic bone compared to the control bone in the mid-cortex (Figure 6b). Similar to the results of the cluster nodes, we found that there is a low proportion of hubs within the LCN, but there is a 39% (p=0.17) increase in the number of hubs in the diabetic bone from the mid-cortex (Figure 6c).

**Figure 6:**
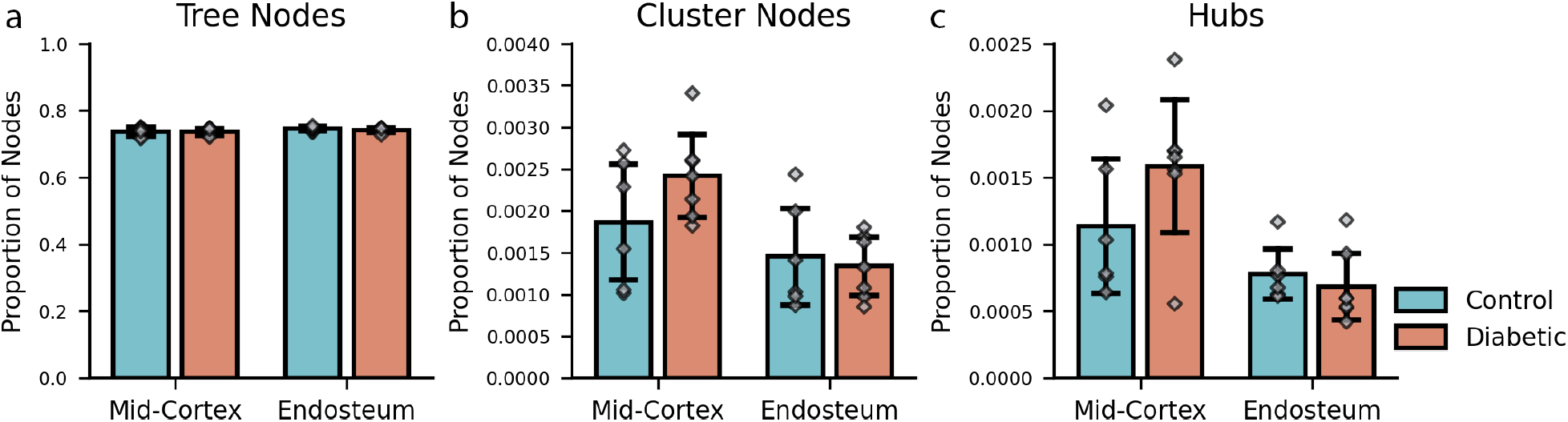
There is an elevated proportion of high-degree nodes in the mid-cortex of diabetic bone. (**a**) A bar graph showing the proportion of tree nodes for each group. Nodes are classified as tree nodes if they are of degree 3 and have a clustering coefficient (CC) of 0. For all groups, the majority of the nodes in the network were tree nodes. (**b**) A bar graph showing the proportion of cluster nodes for each group. Nodes are classified as cluster nodes by having a CC greater than 0.05. (**c**) A bar graph showing the proportion of hubs for each group. Nodes are determined hubs if they have a degree greater than or equal to 8. Note that some nodes are included in both the hubs and cluster node proportions. All bar graphs show the mean and standard deviation for each measurement.

## Discussion

To investigate the effect of type 2 diabetes on the microstructure of cortical bone, we imaged the LCN of samples from control and diabetic (ZDSD) rats. We acquired 3D images of the LCN using CLSM, segmented the LCN using deep learning, and converted the segmentation into a topological graph of edges and nodes. Using this representation of the LCN, we used connectomic approaches to investigate the impacts of diabetes on the structure and organization of the LCN cortical bone. Our results indicate that type 2 diabetes significantly impacts the LCN in cortical bone. The impacts of diabetes are observed in the mid-cortex of the cortical bone but not near the endosteum. In the mid-cortex, we found that diabetes significantly reduces the density of the LCN through a 21% reduction in canalicular density and a 30% reduction in node density. As a result of the decreased LCN density, we found that in the mid-cortex, diabetes reduced radial permeability by 41%. Additionally, due to the reduced LCN density in the mid-cortex, we found that diabetes caused the bone matrix to be located 18% further from the LCN. We also observed that diabetes causes a slight increase in cluster nodes and hubs within the network compared to the control bone in the mid-cortex.

A key impact that diabetes had on the bone was the reduction in canalicular and node density of the LCN within the mid-cortex. These reductions in network density could indicate reduced osteocyte function in this region of bone. Other groups have observed a similar reduction in overall LCN density due to aging [15, 24, 26]. Schurman et al. found that reduced canalicular density contributed to diminished diffusivity and that reductions in canaliculi compromised key osteocyte functions, such as mass transport and mechanosensation [26]. Similarly, the present study found that reduced canalicular density was accompanied by reduced LCN permeability. This reduced permeability could negatively impact bone quality and bone maintenance by limiting osteocyte functionality. Diabetes significantly impacts the LCN in the mid-cortex, which shows evidence of reduced functionality and mass transport.

The reduced canalicular density also causes the bone matrix to be situated further from the LCN. In the mid-cortex, where we observed reduced canalicular density with diabetes, we also observed that the bone matrix was further from the LCN. With more bone matrix situated further away from the LCN, mechanosensation, and remodeling rate might be decreased, diminishing bone quality. One example of how this reduced canalicular density could impact remodeling is how it could impact perilacunar canalicular remodeling (PLR), which relies on the acidification of its surroundings [31]. Lower canalicular density would result in less bone being in a region that would be reached by this acidification during PLR. This provides another example of how diabetes may impact the LCN in the mid-cortex, which could reduce bone quality, functionality, and maintenance.

Diabetes was previously shown to decrease bone’s resistance to fracture in the same ZDSD rat model [19]. Woolley et al. analyzed lacunar density, volume, and whole-bone mechanical properties of diabetic and control rat femurs. They attributed decreases in fracture resistance in diabetic rat femurs to increases in AGEs and changes in bone microstructure, namely osteocyte lacunae [19]. The present study identifies further evidence that diabetes deteriorates bone’s microstructure. In the mid-cortex, we observed that diabetes reduced canalicular density and increased hub proportion (Figure 3, Figure 6). These alterations to the LCN could negatively impact its functions, such as communication and remodeling because the LCN density is reduced. Reductions in LCN remodeling could cause the bone to be more susceptible to fracture [14, 32].

Osteocytes locally remodel their surroundings, an important function to maintain bone quality [14]. We gain insight into osteocytes and their function by analyzing their lacunae with 3D imaging techniques, such as CLSM or micro-computed tomography. Using the CLSM images, we observed a 43% increase in lacunar density in the mid-cortex and no change in lacunar density near the endosteum (Figure 3). These results contrast with the results of Woolley et al. They saw decreased lacunar density in diabetic rat femurs compared to controls when analyzing these metrics for the entire cross-section of the bone using micro-computed tomography [19]. We expect the discrepancy between these studies is due to the regional dependence of lacunar density [28]. Diabetes may cause a regional increase in lacunar density in the mid-cortex of the anterior medial quadrant of the bone and cause a general decrease in lacunar density throughout the entire cross-section. Other researchers have also observed an increased lacunar density with diabetes using CLSM [17]. The present study’s findings further support the concept that the impacts of diabetes on bone may occur in specific regions.

We found that the canaliculi are more aligned near the endosteum than the mid-cortex. This difference was observed regardless of whether the bone came from the control or diabetic rat group. In the endosteum, the canaliculi were generally dense and highly aligned in a primary direction, while in the mid-cortex, the canaliculi were less aligned and adopted more of a radial organization (Figure 1). These results agree with what other researchers have reported [28, 29]. Kerschnitzki et al. reported that cortical murine bone has organized canaliculi in the bone’s endosteal and periosteal regions, consisting of lamellar bone [29]. They also observed more disorganized canaliculi in the mid-cortex or microlamellar bone. Shipov et al. confirmed that this organizational variation is present in cortical rat bone [28]. This regional variation in morphology explains the marked organizational changes we see comparing the aligned canaliculi near the endosteum to the radial canaliculi of the mid-cortex. However, these regional variations do not account for the trend toward increased mean canalicular angle observed in the mid-cortex of diabetic rat femurs (Figure 1e). This change in orientation angle indicates that diabetes impacts the organization of canaliculi in the mid-cortex. Organizational changes of the LCN have been connected to other properties of bone architecture, such as collagen and mineral orientation [29, 33]. Kerschnitzki et al. found that canaliculi are generally oriented perpendicular to the collagen matrix [29] and that mineral particle orientation and thickness correspond to the canalicular organization. They found that organized regions of canaliculi have well-oriented mineral particles [33]. These findings by Kerschnitzki et al. show that canalicular organization can be used as a proxy to estimate collagen and mineral orientation. Variations in organization highlight that bone quality varies throughout different regions of the bone and emphasize the importance of consistently selecting regions within the bone to compare LCN network between different groups.

Another impact of diabetes on the LCN was a modest increase of highly connected junctions, or hubs, in the network in the mid-cortex. In this study, most of the analyzed samples’ nodes were classified as tree nodes (degree of 3 and CC of 0). These nodes are advantageous if a disruption, such as a microcrack, occurs because it would not impact a large portion of the network [17, 33]. A small proportion of highly connected hubs was present in all samples. These hubs are a noteworthy feature of the LCN; if these highly connected junctions are destroyed, the LCN could be disrupted [17, 30]. Regionally, the proportion of cluster nodes and hubs is higher in the mid-cortex compared to the endosteum in both the control and diabetic groups. This increase could be attributed to the radial organization in the mid-cortex, which may lend itself to higher degree connections than the parallel aligned endosteum. A moderate increase of these hubs was observed within the mid-cortex with diabetes. This increase in the proportion of hubs in diabetic bone has been observed in the literature. Mabilleau et al. observed a significant increase in hubs associated with diabetes in high-fat diet conditions. They explain that these hubs may indicate new dendritic connections established during remodeling are likely to occur at already highly connected regions [17]. This increase in highly connected regions in the LCN of diabetic bone alters the organization and communication flow within the network and increases the risk of disruption if a microcrack disrupts this portion of the network [17].

Some limitations accompany the findings of this study. A key limitation of this study is that the imaging technique labeled all voids within the bone and not the cellular components. This resulted in analyzing lacunae and canaliculi rather than osteocytes and dendrites [34]. Diabetes has been shown to increase osteocyte apoptosis, resulting in more empty lacunae [2, 35, 36]; however, our imaging approach did not allow us to distinguish empty lacunae. Analyzing the canaliculi instead of the dendrites likely overestimates the number of dendrites. Tiede et al. found that the number of canaliculi is 1.2–1.7 fold higher than the number of dendrites [15]. In future work, it would be beneficial to stain the LCN and the cellular components using a multiplexed imaging protocol [34]. This approach would allow for the analysis of empty lacunae and further describe the impact of diabetes on osteocyte functions. Another imaging-related limitation is the small field of view of confocal microscopy. We sample two 184×184×42 µm regions within the cross-section of the femur to evaluate the impacts of diabetes on the LCN. This results in the majority of the cross-section not being analyzed. It would be interesting to sample more regions throughout the bone to quantify the effect of diabetes in additional regions. Although we only analyzed two regions within the bone, our consistent selection of these regions allowed us to detect regional changes in the bones’ microstructure with diabetes. The connectomic analysis of the LCN also has some limitations. One analysis limitation of this study is the evaluation of the node degree. This measurement is sensitive to the approach selected to merge nodes. Selecting different merging algorithms or neighborhood radii could produce different results. In this study, we selected the length of the 3D diagonal of a voxel as the neighborhood radii to merge the nodes.

In conclusion, type 2 diabetes significantly impacts the LCN in ZDSD rat femurs. We observed evidence of this impact in reductions of canalicular and node density within the LCN. These reductions could reduce bone quality, functionality, and maintenance. We observed these impacts in the mid-cortex of the femur but not in the bone near the endosteum. This study adds to the growing research on the impact of type 2 diabetes on bone. Specifically, we show that diabetes reduces the properties of the LCN in specific regions within the bone. These results highlight that the impacts of diabetes are not universal throughout the bone.

## Methods

### Animals and tissues

Sprague Dawley rats were used to model the effects of diabetes on the LCN. The specific rat model utilized was the Zucker Diabetic Sprague Dawley (ZDSD) rats (Diabetic, N=7), which were compared to lean Sprague Dawley rats (Control, N=6). All rats were male and purchased from Charles River. Details on this model can be found in the papers by Reinwald et al. [37] and Wooley et al. [19]. Briefly, the rats were housed and treated following protocols approved by the Institutional Animal Care and Use Committee (IACUC). The rats were fed standard chow (Catalog No. 2920X; Envigo Teklad Global). All rats were euthanized at 19 weeks, and the left femora were harvested from the rats. Transverse sections were cut from the mid-diaphysis of the femur using a bathed low-speed saw. The anterior medial quadrant of the section was isolated to evaluate bone that would experience compression.

### Bone staining

The transverse sections of bone were polished to 200 µm using an electric polishing table and then polished by hand to 130 µm. The samples were sonicated for 15 minutes to remove debris that could obstruct the stain from fully infiltrating the bone. The samples were dehydrated in graded ethanol baths of 50%, 75%, 90%, and 100% for ten minutes each. The samples were stored in 100% ethanol until staining.

The samples were stained with fluorescein isothiocyanate (FITC, Sigma Aldrich F7250), which stains the walls of the void space within the bone. The stain was prepared by combining FITC with ethanol (1mg/100mL). The stain was mixed on a stir plate for one hour and passed through a 0.02 µm filter. Each bone sample was submerged in 20 mL of stain and placed on a rocker for four hours to allow the stain to permeate the bone. After staining, the samples were rinsed in 100% ethanol for one hour and then further polished to 100 µm. The purpose of the final polishing step is to remove any unwanted stain that may have adhered to the surfaces of the bone. The bones were sonicated for 15 minutes and then rinsed in 100% ethanol for another hour. The bone samples underwent tissue clearing by placing them in a solution of one part benzyl alcohol and two parts benzyl benzoate (BABB) [38] overnight. The following day, the samples were rehydrated in graded PBS baths of 50%, 75%, 90%, and 100% for ten minutes each. The samples were mounted on a glass slide with a mounting medium to prevent photobleaching (EMS Glycerol Mounting Medium with DABCO cat. 17989-50).

### Confocal microscopy

The stained samples were imaged using a Leica SP8 White light Laser Confocal microscope with a 63x objective (HC PL APO CS2 63x/1.40 OIL) and a photomultiplier tube detector (PMT). The 1024×1024 image format was used with a zoom of 1, resulting in a voxel spacing of 0.18×0.18×0.3 µm. The final imaged volume was 184×184×75 µm. Images were acquired in two regions of each bone sample, one in the well-organized regions near the endosteum and another in the disorganized mid-cortex [28, 29].

### Image processing and segmentation

The analysis in this study required the confocal images to be segmented. Each voxel in the image was assigned to either background, canaliculi, lacunae, or blood vessel. To prepare the confocal image stacks for segmentation, the images’ histograms were equalized using clipped local histogram equalization (CLAHE) and denoised using a 3D Gaussian filter. A subset of each image stack was segmented using tools in Dragonfly to provide training data to the U-net to learn to perform the segmentation task [39]. The segmented images were divided into 256×256 patches for training the U-net.

A U-net was trained to perform the segmentation task because of its capability to label the different features despite the noise and common grayscale values between features. The U-net was trained in PyTorch [40] using cross-entropy as the loss function and Adam as the optimizer [41] with an initial learning rate of 0.0001. A learning rate scheduler reduced the learning rate by half every 20 epochs. The training data was passed to the network in mini-batches consisting of 32 images. The images were randomly flipped vertically and horizontally to augment the data. The network was trained for 100 epochs, with the condition that training would stop after 10 epochs with no improvement in the validation data. The model weights were saved after each epoch with a reduction in validation loss. The U-net reached the stop criteria after 97 epochs, at which point the best model weights were loaded back into the model. This trained U-net was used to segment all of the image stacks. Each segmented image was reviewed, and manual corrections were performed as needed.

### Canaliculi alignment and spacing

The segmented images were used to determine the alignment of the canaliculi. The canaliculi were isolated using the segmentation. To minimize the impacts of lacunae and blood vessels on the alignment, the segmented lacunae and blood vessels were dilated in 3D and then subtracted from the canaliculi segmentation using Fiji [42]. The 2D alignment of the remaining canaliculi was determined using the OrientationJ plugin in Fiji [42, 43]. For each sample, the distribution of orientations was averaged for all slices to obtain a single distribution per sample.

Canalicular spacing was measured using the Fiji plugin BoneJ [42, 44]. The segmented canaliculi, lacunae, and blood vessels were considered foreground, and the trabecular spacing function in BoneJ was used to estimate the canalicular spacing.

### Network extraction

A base graph representing the canaliculi structure, the lacunae, and blood vessels was computed for each labeled sample. First, a signed distance field to the foreground/background interface was generated, with negative values inside the foreground. The topological 1-skeleton was extracted as the minimum-1-saddle arcs of the Morse-Smale complex using MSCEER [45, 46]. This complete graph connecting all valley-like structures was simplified using an approach that preserved the topological structure of the foreground [47]. The topological 1-skeleton might have contained overlapping lines, which were corrected by introducing nodes at each junction point and merging overlapping lines [48, 49]. The graph was cut at the foreground/background interface, keeping only the foreground. A user-specified length was used to iteratively prune-and-merge short leaves and degree-2 nodes from the graph [50]. Finally, nodes were inserted along edges at the interfaces between canaliculi, lacunae, and blood vessels.

### Network analysis

Using the graph extracted from the segmented image stack, we determined connectomic metrics, such as canalicular density, node density, and node classification [25, 30]. Canalicular density was determined by determining the total canalicular length from the graph and dividing that length by the scan volume. The node classification was performed similarly to Kollmannsberger et al. [30]. However, the node degree and classification results by Kollmannsberger et al. [30] are not directly comparable to our graph since their graph computation implicitly iteratively merges nearby nodes using a 26-neighborhood. To obtain comparable results for node degree, we virtually collapse all edges shorter than the 3D diagonal of a voxel (only graph connectivity matters, not geometry, in counting node degree). Following the virtual collapse of edges, we used Networkx [51] to determine the CC for each node. The CC measures the connectivity of the immediate neighboring nodes. This metric ranges from 0 to 1, where 0 indicates the neighboring nodes are unconnected and 1 indicates all possible connections are formed. We classify nodes as tree and cluster nodes. Tree nodes have a degree of 3 and a CC of 0. Cluster nodes have any degree and a CC greater than 0.5 [30].

### Permeability computation

We directly compute the directional permeability of samples using the lacunae-canaliculi network (LCN) as a pore-network model (PNM) [52]. Unlike Kozeny permeability estimation using porosity and pore surface area, this network-based approach accounts for anisotropy and the presence of dead-ends within the structure. In the PNM, nodes of the LCN represent junctions where canaliculi meet or where they connect to lacunae, while edges correspond to either individual canaliculi or internal pathways within lacunae. Where an edge of the LCN crosses between pixels labeled canclicul or lacunae (by the ML classifier), additional degree-2 nodes are created to split edges such that each resulting edge is entirely canaliculi or lacunae.

The overall approach computes the directionaly permeability of a sample is by applying Darcy’s law,

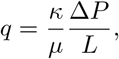

where *q* is the volumetric flow rate, *κ* is the permeability, *µ* is the dynamic viscosity, Δ*P* is the pressure drop, and *L* is the length of the domain. In the LCN, fluid flow induced by stress-strain is restricted to the pore network. Instead of using a full computational fluid dynamics (CFD) simulation, a pore-network model (PNM) provides a reasonable proxy for estimating the permeability and fluid flow behavior within the network.

In this PNM, flow is computed along *pores* between *reserviors*; LCN nodes become reservoirs, and LCN edges pores. The nodes of the LCN touching the domain boundaries in the three axial directions become inlets/outlets, and edges are assigned conductances *G* are computed using the Hagen–Poiseuille equation,

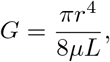

with *r* and *L* denoting the radius and length of each pore. Due to limitations in image resolution, rather than attempting to compute the individual radii of canaliculi and lacunae, we uniformly assign either the mean canalicular radius or the mean lacunar radius to a pore, depending on the segmentation value.

A pressure difference applied across the sample induces the fluid flow through the network. This formulation is practically solved by translating it into a resistor network model, where a matrix is used to represent the system of linear equations derived from Kirchhoff’s Current Law (KCL) and the conductance relationships between nodes. The network is augmented with edges having 1*/ϵ* conductance connecting the inlets with a virtual node for injecting current and connecting the outlets with a node representing virtual ground. The system is represented as *AV* = *B*, where *A* is the sparse *contuctance matrix, V* is the vector of unknown node voltages, and *B* is the vector representing net currents injected at each node. Solving for *V* gives not only the overall conductance (from the voltage difference between the virtual inlet node and virtual ground node, and the injected current), but also the current carried by each edge of the network. The PNM solver was implemented using `spsolve` in `scipy`.

Subdomains were selected from each image stack to ensure the network included distinct organizations from the mid-cortex and endosteum. This was achieved by selecting specific regions of the image stacks. For the mid-cortex volumes, the region furthest from the endosteum was selected. For the endosteal volumes, the region closest to the endosteum was selected, ensuring that the bone marrow cavity wasn’t included. These subdomains had a volume of 61×184×42 µm. To normalize the directional permeability of the sample, we use

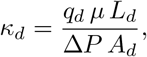

where *κ*_*d*_ is the permeability in the direction *d* (*x, y*, or *z*), *q*_*d*_ is the volumetric flow rate (equivalent to the total current in the resistor network) in the direction *d, µ* is the dynamic viscosity of the fluid, *L*_*d*_ is the length of the sample along the flow direction, Δ*P* is the pressure difference (computed voltage) across the sample, and *A*_*d*_ is the cross-sectional area perpendicular to the flow direction.

### Statistics

The control and diabetic groups were compared using a student t-test from Scipy [53]. These groups were compared in both the bone’s mid-cortex and endosteal regions. Differences were considered significant if the p-value was less than 0.05.

## Acknowledgements

This work received support from the National Science Foundation under NSF CAREER Grant CMMI 2045363 and the Washington University Diabetes Research Center (NIH NIDDK P30 DK020579). This work was partly supported by the National Institute for Occupational Safety & Health (NIOSH) #5T42OH008414-16. The opinions and findings of the authors do not necessarily reflect the views and opinions of NIOSH. We acknowledge the Cell Imaging Core at the University of Utah for using the Leica SP8 White light Laser Confocal microscope and thank Michael Bridge for assistance in image acquisition.

## Author Contributions

C.A., M.S., A.G., and V.P. conceived and designed the experiments. M.S. and I.J. prepared, stained, and imaged the bone samples. M.S. and I.J. completed the image segmentation. A.G. and M.S. performed the network analysis. M.S., C.A., A.G., and V.P. interpreted the data. M.S., C.A., and A.G. wrote the manuscript with contributions from all coauthors.

## Data Availability

The confocal image and segmentation dataset used in this publication are available in this Zenodo repository https://doi.org/10.5281/zenodo.11061867[54].

## Notes

### Competing Interest Statement

The authors have declared no competing interest.

